# Lipid-specific labeling of enveloped viruses with quantum dots for single-virus tracking

**DOI:** 10.1101/2020.01.06.895805

**Authors:** Li-Juan Zhang, Shaobo Wang, Li Xia, Cheng Lv, Hong-Wu Tang, Gengfu Xiao, Dai-Wen Pang

**Affiliations:** Key Laboratory of Analytical Chemistry for Biology and Medicine (Ministry of Education), College of Chemistry and Molecular Sciences, State Key Laboratory of Virology, The Institute for Advanced Studies, and Wuhan Institute of Biotechnology, Wuhan University, Wuhan, People’s Republic of China; Wuhan Institute of Virology, Chinese Academy of Sciences, Wuhan, People’s Republic of China; State Key Laboratory of Medicinal Chemical Biology, Tianjin Key Laboratory of Biosensing and Molecular Recognition, Research Center for Analytical Sciences, College of Chemistry, and School of Medicine, Nankai University, Tianjin, People’s Republic of China

**Author notes:** Corresponding Author (D.-W. P.); (G. X.). These authors contributed equally to this work (Li-Juan Zhang, Shaobo Wang).

## Abstract

Quantum dots (QDs) possess optical properties of superbright fluorescence, excellent photostability, narrow emission spectra, and optional colors. Labeled with QDs, single molecules/viruses can be rapidly and continuously imaged for a long time, providing more detailed information than labeled with other fluorophores. While they are widely used to label proteins in single-molecule tracking studies, QDs have rarely been used to study virus infection, mainly due to lack of accepted labeling strategies. Here, we report a general method to mildly and readily label enveloped viruses with QDs. Lipid-biotin conjugates were used to recognize and mark viral lipid membranes, and streptavidin (SA)-QD conjugates were used to light them up. Such a method allowed enveloped viruses to be labeled in 2 hours with specificity and efficiency up to 99% and 98%. The intact morphology and the native infectivity of viruses could be furthest preserved. With the aid of this QD labeling method, we lit wild-type (WT) and mutant Japanese encephalitis virus (JEV) up, tracked their infection in living Vero cells, and found that H144A and Q258A substitutions in the envelope (E) protein didn’t affect the virus intracellular trafficking. The lipid-specific QD labeling method described in this study provides a handy and practical tool to readily “see” the viruses and follow their infection, facilitating the widespread use of single-virus tracking and the uncovering of complex infection mechanisms.

**Author summary:** Virus infection in host cells is a complex process comprising a large number of dynamic molecular events. Single-virus tracking is a versatile technique to study these events. To perform this technique, viruses must be fluorescently labeled to be visible to fluorescence microscopes. Quantum dot is a kind of fluorescent tags that has many unique optical properties. It has been widely used to label proteins in single-molecule tracking studies, but rarely used to study virus infection, mainly due to lack of accepted labeling method. In this study, we developed a lipid-specific method to readily, mildly, specifically, and efficiently label enveloped viruses with quantum dots by recognizing viral envelope lipids with lipid-biotin conjugates and recognizing these lipid-biotin conjugates with streptavidin-quantum dot conjugates. Such a method is superior to the commonly used DiD/DiO labeling and the other QD labeling methods. It is not only applicable to normal viruses, but also competent to label the key protein-mutated viruses and the inactivated high virulent viruses, providing a powerful tool for single-virus tracking.

## Introduction

Single-particle tracking is a powerful tool to study the dynamic molecular events in living cells. An essential prerequisite to perform this technique is fluorescently labeling the targets. In the past decade, various fluorescent tags such as organic dyes [1], fluorescent proteins [2], metal complex of dppz [3], and QDs [4] have been used to label the target molecules/viruses. The excellent optical properties make QDs unparalleled in single-molecule/virus tracking. Single molecules/viruses illuminated with QDs can be rapidly and continuously tracked for a long time [5, 6], and their interactions with multiple other molecules can be monitored simultaneously [7-9], providing more detailed information to dissect cellular events than those labeled by other fluorophores. Thanks to these advantages, QDs have been widely used to label proteins for single-molecule tracking studies [10-17]. But due to lack of accepted labeling method, QDs were rarely used to label viruses, which in turn limited the widespread use of single-virus tracking.

To label viruses with QDs, more than a dozen of methods have been developed, which could be roughly divided into three groups. By directly (*e.g.*, virus-NH_2_-COOH-QD) or indirectly (*e.g.*, virus-NH_2_-COOH-biotin-SA-NH_2_-COOH-QD) attaching QDs to the amino on viral proteins, both enveloped and non-enveloped viruses can be labeled (*group 1*) [18, 19]. Similarly and more ingeniously, QD-labeled viruses can be obtained by genetically engineering specific viral proteins to combine them with reactive biomolecules and then the correspondingly modified QDs (*group 2*) [20, 21]. Besides, by modifying the membrane of host cells and propagating viruses in them, viruses with reactive membranes can be harvested and then labeled with QDs (*group 3*) [22, 23]. Although so many methods reported, none of them has been broadly used in practical studies due to the concerns that they may affect the bioactivity of the target proteins (*group 1*), they are too complicated and time-consuming (*group 2*), or the labeling efficiency greatly varies with the cell and the virus (*group 3*).

The aim of this work is to provide a universal and convenient method to specifically and efficiently label enveloped viruses with QDs while preserving the native state of viral proteins. In conventional virology, lipophilic dyes such as DiO and DiD that can readily insert into lipid bilayer membranes are widely used to label viruses. Learning from these long-chain lipophilic dyes, we developed a convenient method to label viruses with QDs by modifying viral lipid membranes with lipid-biotin conjugates and lighting these extraneous lipids up with SA-QD conjugates. Such a method could leave viral proteins uninvolved, and its effect on viral infectivity was negligible. It allowed enveloped JEV, porcine reproductive and respiratory syndrome virus (PRRSV), and influenza A virus (IAV) to be labeled with specificity and efficiency above 95% and 93%, respectively. The whole labeling procedure comprised just five brief steps and could be done within 2 hours. With the aid of this lipid-specific QD labeling method, both WT and E protein-mutated JEV were fluorescently labeled, and their infection behaviors were thus visually analyzed.

## Results and discussion

### Labeling design

Labeling with high specificity and high efficiency and without affecting virus infectivity is essential to obtain hi-fi information about virus infection, while labeling with great convenience and universal applicability is essential for a method to be widely used. To develop a QD labeling method meeting these requirements, we learned from lipophilic dyes and designed a strategy to label viruses by targeting the lipid membrane. An amphipathic lipid-biotin conjugate, DSPE-PEG-Biotin (Fig 1A), was used to recognize viral lipid membranes and mark them with biotin, and SA-QD conjugates were used to combine with the exogenous lipid through interaction with biotin and thus light the virus up (Fig 1B). As seen in S1 Fig, DSPE-PEG-Biotin could insert into lipid membranes as fast as DiD. After incubation with DSPE-PEG-Biotin for 30 min and then with SA-QD for 10 min, cells could be efficiently labeled with QDs. To apply this strategy to viruses, we optimized the labeling procedure as illustrated in Fig 1C: clearing cell debris from virus solution by low-speed centrifugation and syringe filtration, biotinylating viral lipid membranes by incubation with DSPE-PEG-Biotin under shaking, removing unincorporated lipid-biotin molecules by gel filtration, pre-attaching biotinylated viruses to cell surfaces by incubation with cells at 4°C, and coupling SA-QDs to the lipid-biotin on viral membranes by incubation with the cells at 4°C. Unbound viruses and QDs could be removed just by washing the cells. Such a strategy can thoroughly evade ultracentrifugation, dialysis, and ultrafiltration processes that are indispensable for removing the cell-derived reactive molecules, redundant functional reagents, unlabeled viruses, or unbound QDs in many other labeling strategies [24-30]. This strategy furthest minimized and simplified the handling of viruses, making the QD labeling milder and more convenient.

**Fig 1.**
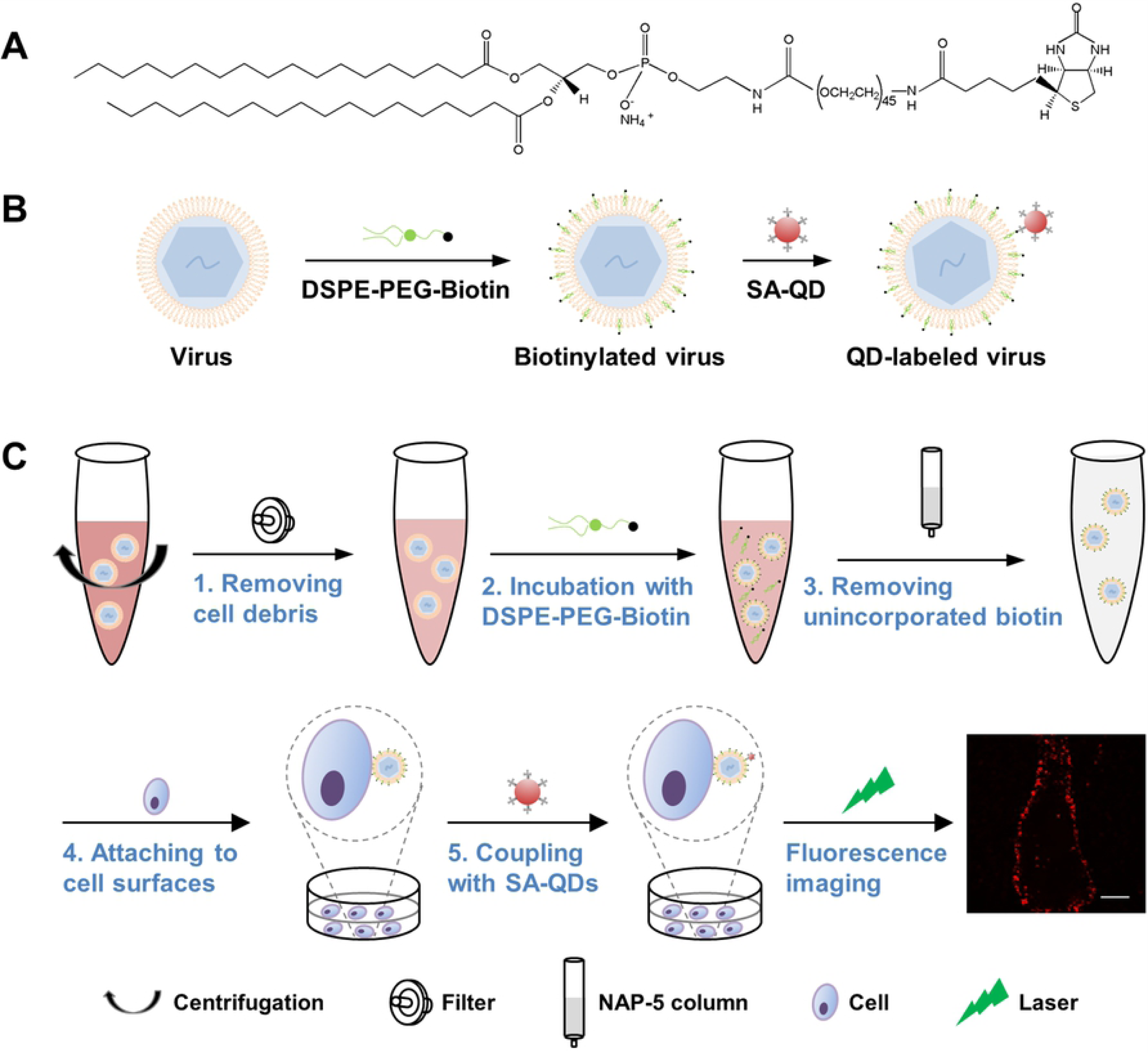
Lipid-specific QD labeling of enveloped viruses. (A) Structure of DSPE-PEG (2000)-Biotin. (B) Using the rapid insertion of the lipid-biotin conjugate into lipid membranes and the specific high-affinity interaction between biotin and SA to label viruses. (C) The entire labeling procedure comprising five brief steps (*1–5*). The last panel is a fluorescence image of JEV labeled as thus on a Vero cell. Scale bar, 10 μm.

### Specifically, efficiently, and mildly labeling viruses

JEV about 50 nm in diameter was used as the model virus to experimentally evaluate the labeling strategy. Raw JEV and biotinylated JEV were prebound to glass slides, respectively, and labeled with SA-QD 705 and anti-E protein-DyLight 488. As seen in Fig 2A, there was no obvious QD signal colocalized with DyLight-stained raw JEV, while almost all the DyLight-stained biotinylated JEV was colocalized with QDs. These data indicated that DSPE-PEG-Biotin could insert into the lipid membrane of viruses, and SA-QDs could efficiently bind to viruses modified with the lipid-biotin conjugate specifically through interaction with biotin. The overlapped fluorescence peaks of QD and DyLight (Fig 2B) and the scarcely any negative values of the product of differences from the mean (PDM) of pixel intensities in the two channels (Fig 2C) visually showed that almost all the QD and DyLight signals were colocalized. Statistically, about 99% QD signals were colocalized with the DyLight-stained viruses (tM_QD_ = 0.986 ± 0.008), and about 98% viruses were colocalized with QDs (tM_DyLight_ = 0.976 ± 0.021) (Fig 2D). In other words, the QD labeling specificity and efficiency on glass slides were 99% and 98%, respectively. The high intensity correlation quotient (ICQ) value (0.298 ± 0.014) confirmed this nearly complete colocalization further [31]. Labeling viruses on Vero cell surfaces showed that the QD and DyLight signals still colocalized to a very high degree (S2A–C Figs). The tM_QD,_ tM_DyLight,_ and ICQ values were 0.979 (± 0.018), 0.957 (± 0.030), and 0.291 (± 0.026), respectively (S2D Fig). The specificity and efficiency of this method are superior to those of the previously reported QD labeling methods to different degrees and significantly superior to the specificity and efficiency of DiD and DiO labeling (S3 Fig).

**Fig 2.**
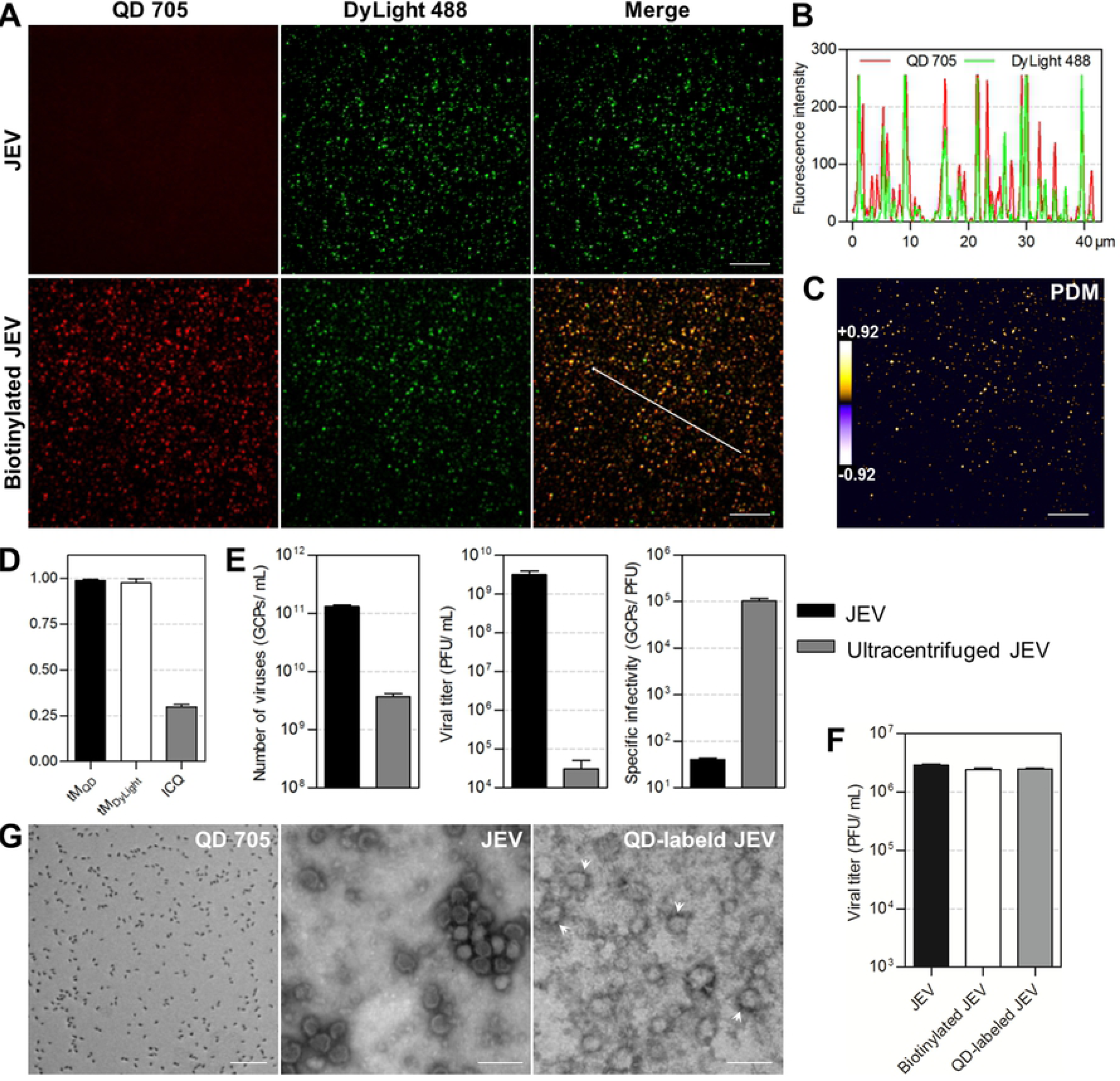
Specifically, efficiently, and mildly labeling JEV with QDs. (A) JEV and biotinylated JEV were prebound to glass slides and labeled with SA-QD 705 (red) and anti-E-DyLight 488 (green). (B) Line profile showing distributions of the signals on the line in A. (C) PDM image showing the colocalized (PDM > 0) and uncolocalized (PDM < 0) spots in the lower merge panel in A. Scale bars, 10 μm. (D) The tM_QD_, tM_DyLight_, and ICQ values calculated from 20,000 viral particles from three experiments. (E) The number of genome-containing particles (GCPs), titers, and specific infectivity of viruses before and after ultracentrifugation. (F) Titers of viruses before and after biotinylation and SA-QD 705 labeling (n = 3). (G) TEM images of SA-QD 705, JEV, and QD-labeled JEV (arrowheads). Scale bars, 100 nm.

To determine the effect of QD labeling on viruses, both the pretreatment and the labeling processes were analyzed. In our lipid-specific method, viruses were just processed with low-speed centrifugation and syringe filtration before labeling, while in many other methods, they would need further purification by ultracentrifugation [32, 33]. Comparing un-ultracentrifuged viruses with viruses ultracentrifuged under the generally used conditions showed that high-speed centrifugation greatly reduced virus infectivity (Fig 2E). By evading such violent pretreatment, the native infectivity of viruses was greatly preserved. During the labeling process, no cumbersome operation was performed, and no interaction involving viral proteins was used. Measuring the titer of viruses before and after QD labeling showed that the labeling process had no obvious effect on virus infectivity (Fig 2F). As seen in the transmission electron microscope (TEM) image, QD-labeled viruses were morphologically as intact as unlabeled viruses (Fig 2G). In aggregate, labeling viruses with QDs by the above lipid-specific method could preserve virus infectivity furthest.

### Stably and universally labeling viruses

Under the labeling conditions we used, about 2836 DSPE-PEG-Biotin molecules were incorporated into the lipid membrane of JEV during biotinylation (S4 Fig), and 2 or 3 QDs were coupled to the biotinylated virus afterwards (S5 Fig). To evaluate the stability of QD combining with viruses, we dually labeled JEV with QD 605 and QD 705 and allowed the viruses to infect Vero cells for different time. It could be observed that the two kinds of QDs kept colocalized with the DyLight-stained viral envelope in 2 hours of virus infection (Fig 3A). Almost no QD signal could be observed alone (Fig 3B). The steady Mander’s coefficients and ICQ values of DyLight *vs.* QD 605, DyLight *vs.* QD 705, and QD 605 *vs.* QD 705 suggested that the colocalization relationships among DyLight, QD 605, and QD 705 barely changed during virus infection (Fig 3C). These results indicated that QDs coupled to viruses would not separate from the envelope and could stably point viruses out during virus infection, ensuring getting reliable information.

**Fig 3.**
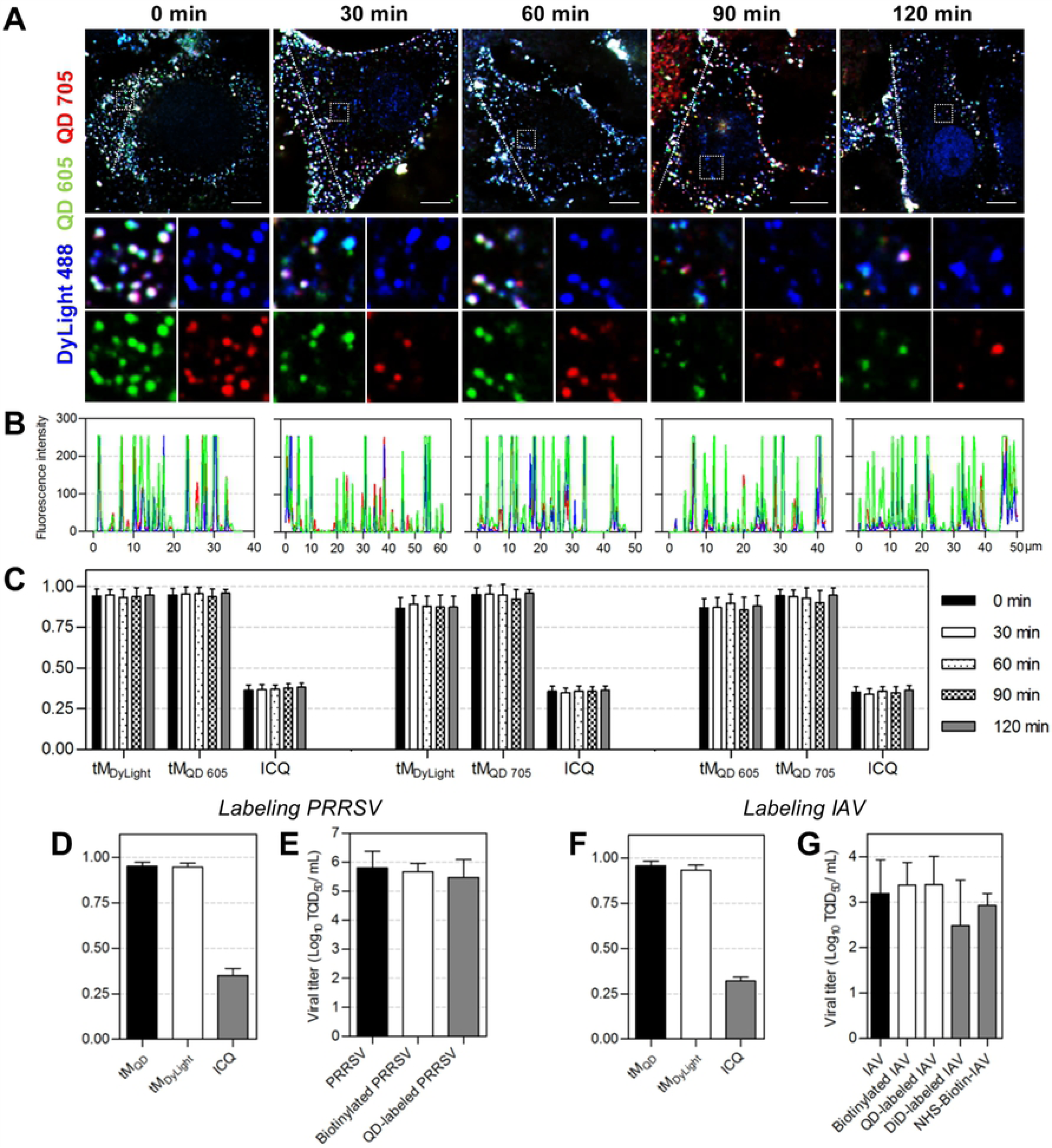
Stability and universality of the QD labeling method. (A) JEV was dually labeled with SA-QD 605 (green) and SA-QD 705 (red). Vero cells infected by the double-labeled viruses for 0, 30, 60, 90, and 120 min were fixed and stained with anti-E-DyLight 488 (blue). Scale bars, 10 μm. (B) Line profiles showing distributions of the fluorescence signals on the lines in A. (C) The tM_DyLight_/tM_QD 605_/ICQ, tM_DyLight_/tM_QD 705_/ICQ, and tM_QD 605_/tM_QD 705_/ICQ values calculated from 30 randomly selected cells. (D–G) PRRSV and IAV were labeled with QDs using the lipid-specific method. D and F are the tM_QD_, tM_DyLight_, and ICQ values calculated from 30 cells. E and G are titers of viruses, biotinylated viruses, QD-labeled viruses, DiD-labeled viruses, and viruses covalently biotinylated with NHS-biotin (n = 3 for PRRSV and 5 for IAV).

Then, we applied the above method to PRRSV and IAV to see how it performed when used to label other enveloped viruses. It was found that almost all the QD and DyLight used to label PRRSV were colocalized with each other with tM_QD_, tM_DyLight_, and ICQ values of 0.950 (± 0.022), 0.946 (± 0.022), and 0.352 (± 0.037), respectively (Fig 3D and S6 Fig). The infectious titers of biotinylated PRRSV and QD-labeled PRRSV were nearly the same as that of the raw PRRSV (Fig 3E), suggesting that QD labeling wouldn’t affect PRRSV infection. When used to label IAV, the method still showed high specificity and efficiency (tM_QD_ = 0.955 ± 0.028, tM_DyLight_ = 0.933 ± 0.027, and ICQ = 0.320 ± 0.022) (Fig 3F and S7 Fig). Comparing with DiD labeling and the QD labeling based on covalent interactions with amino on viral surfaces [19, 34], the lipid-specific QD labeling method showed more superiority in preserving virus infectivity (Fig 3G). These results demonstrated that the method described in Fig 1 was universally applicable for the specific, efficient, and mild labeling of enveloped viruses.

### Imaging the infection of WT and mutant JEV

JEV E protein on the envelope plays essential roles in the virus infection. In our previous work, site mutations have been introduced to the E protein, and several amino acids were proved to be important for the virus membrane fusion [35]. But their roles in virus intracellular transport remain unresolved, since it is difficult to study the dynamic trafficking of viruses by traditional methods. Here we labeled the WT, H144A mutant, and Q258A mutant JEV with QDs to visually analyze the effect of the two substitutions on virus infection. The nearly overlapped one-step growth curves showed that QD labeling had no evident effect on the infectivity of WT, H144A, and Q258A JEV (Fig 4A). To analyze their entry activity, the same amount of WT and mutant viruses were bound to cell membranes (Fig 4B). After virus uptake for different time, cells were transferred to 4°C and viruses remained on cell surfaces were stained with Cy3 to be distinguished from the internalized viruses that was singly labeled with QDs (Fig 4C). By counting the viruses inside cells at indicated time, it was found that after synchronization at 4°C most WT viruses entered cells in the first 25 min, and the number of viruses inside cells plateaued in the next 2 hours (Fig 4D). H144A and Q258A viruses followed similar entry kinetics to the WT viruses. Except for individual time points, the amounts of mutant viruses entered cells at most time points were similar to that of the WT viruses, indicating that replacing the H144 and Q258 amino acids in the E protein with alanine did not affect JEV uptake into Vero cells.

**Fig 4.**
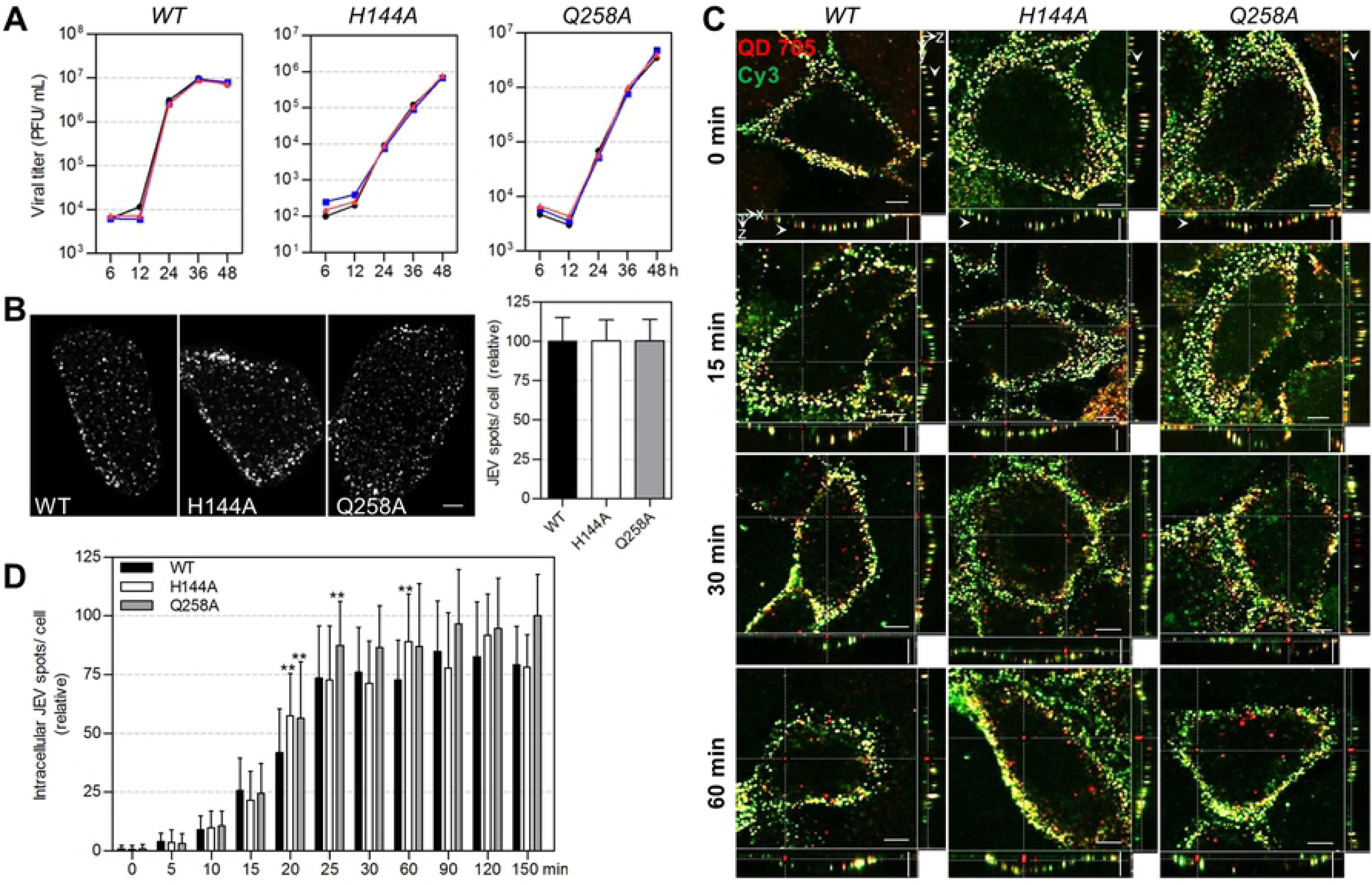
The entry activity of WT, H144A, and Q258A JEV. (A) One-step growth curves of WT, H144A, and Q258A JEV (n=2). The black, blue, and red lines are the curves of raw, biotinylated, and QD-labeled viruses, respectively. (B) WT and mutant JEV were attached to Vero cell surfaces and labeled with QD 705. Cells were imaged in three dimensions (3D) and analyzed with Fiji software. The left panels are the Z-projection images of cells attached with WT, H144A, and Q258A JEV. The histogram is the number of QD-labeled WT/H144A/Q258A JEV on cells (n=100). (C, D) WT and mutant viruses were labeled with QD 705 and allowed to infect cells for 0, 5, 10, 15, 20, 25, 30, 60, 90, 120, and 150 min. Then the cells were transferred to 4°C and incubated with SA-Cy3 for 10 min to stain the viruses remaining on cell surfaces. After fixation, the cells were imaged in 3D and analyzed with Velocity software. C shows cells infected for indicated time. Horizontal and vertical scale bars, 10 μm. D shows the number of viruses internalized in cells after infection for different time (n=30).

Then, we visually analyzed the intracellular transport behaviors of WT and mutant JEV by tracking individual QD-labeled virions. The dynamic transport process of single WT viruses from the cell periphery toward the interior region was observed (Fig 5A and S1 Movie). It could be divided into two stages, viruses moving slowly and irregularly in cell periphery (green lines in Figs 5B–D) and moving rapidly and actively toward the interior of cells (blue lines in Figs 5B–D), according to the speeds, relationships between mean square displacement (MSD) and *Δt* (time interval), and location in cells. As indicated by drug inhibition, the infection of JEV and its rapid active motion in Vero cells were dependent on microtubules and dynein while independent on microfilaments (S8 Fig). Therefore, virus motion in the second stage was the process that dynein drove JEV to move along microtubules toward the interior region. As for the slow irregular motion that differed from the known anomalous or confined motion on cell membranes and slow active motion on microfilaments [36], it was probably the process that JEV diffused across the dense actin-rich region near the plasma membrane. Tracking the movement of single H144A and Q258A JEV virions showed that the two types of mutant viruses moved toward the cell interior in a similar two-stage pattern (Figs 5E–L and S2 and S3 Movies). Statistically analyzing virus speeds in the two stages revealed that H144A and Q258A JEV moved with speeds below 1.0 μm/s in the first stage and with speeds up to several μm/s in the second stage, just as the motion of WT viruses (green and blue histograms in Fig 5M). And the diffusion coefficients and mean velocities of H144A and Q258A viruses in the second stage have no significant differences with those of WT viruses (Fig 5N). These results indicated that these two substitutions in the E protein did not affect the intracellular transport behaviors of JEV.

**Fig 5.**
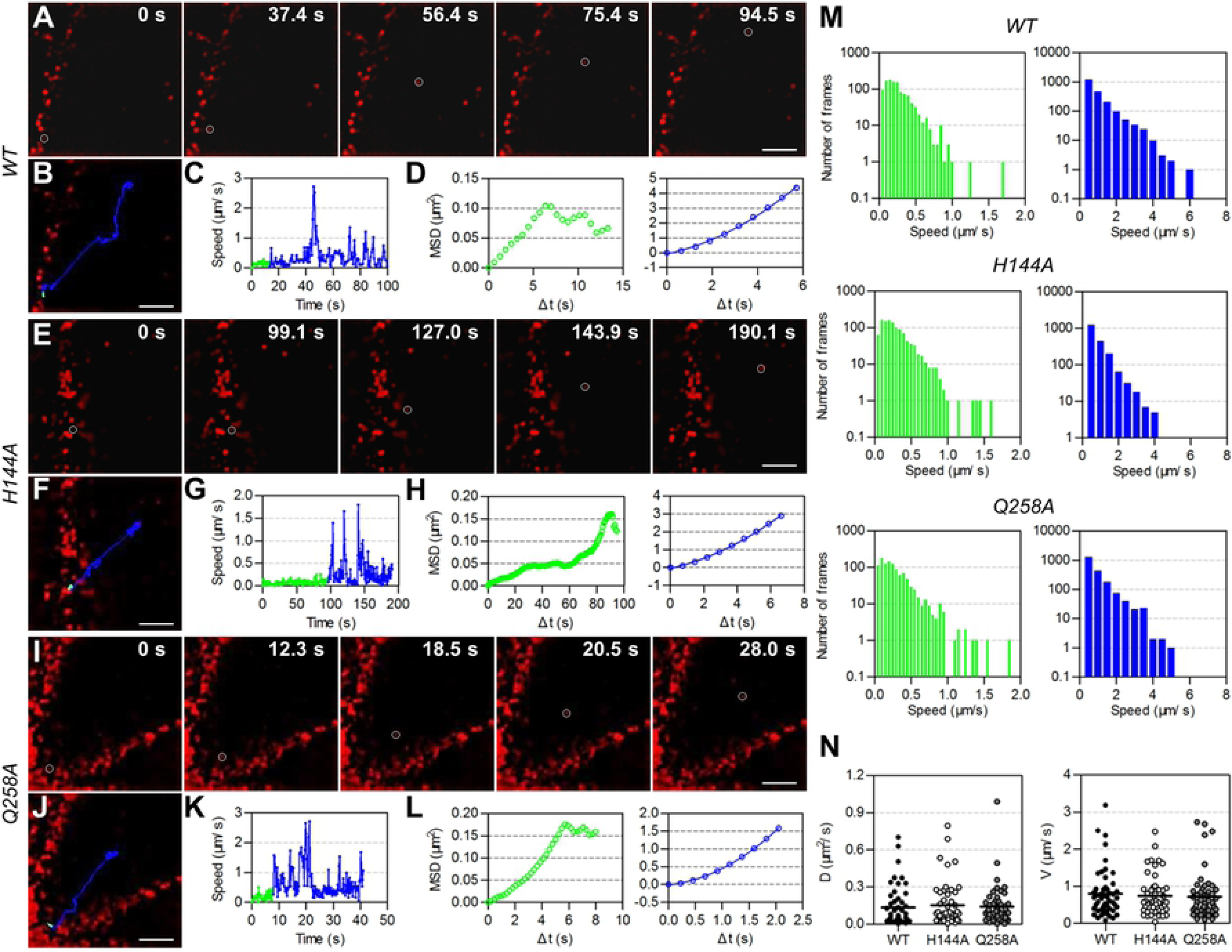
Intracellular transport behaviors of WT, H144A, and Q258A JEV. QD-labeled WT/H144A/Q258A JEV virions were allowed to infect living Vero cells at 37°C and imaged in real-time by a spinning-disk confocal microscope. (A, E, I) Snapshots of QD-labeled viruses (red) infecting cells. (B, F, J) Trajectories of the circled viruses in A, E, and I. (C, G, K) The speed *vs.* time plots of the viruses. (D, H, L) The MSD *vs. Δt* plots of the viruses (green and blue symbols). The green symbols cannot be fitted. The blue lines are the fits to MSD = 4*DΔt* + (*VΔt*)^2^ with *D* = 0.081/0.053/0.039 μm^2^/s and *V* = 0.29/0.19/0.55 μm/s. *D* and *V* are the diffusion coefficient and mean velocity. (M) Statistics of the instantaneous speed of viruses. (N) Statistics of the *D* and *V* of WT/H144A/Q258A JEV moving actively.

Incidentally, we additionally analyzed the fusion activity of the mutant JEV. The viruses were dually labeled with lipophilic DiO and R18 at concentrations allowing R18 to illuminate the viruses consistently and the DiO fluorescence to be quenched before membrane fusion and dequenched after fusion [37]. As thus virus membrane fusion could be determined by measuring the fluorescence intensity of DiO. As seen in S9A Fig, the amount of WT viruses fused with endosomes was greater than that of H144A and Q258A viruses. The fluorescence intensity of DiO in the cells infected by WT viruses increased rapidly in the second and third hours and plateaued gradually in the following 4 hours, while the DiO fluorescence in cells infected by mutant viruses increased very slowly (S9B Fig). The amount of H144A and Q258A viruses fused with endosomes was just 23% and 12% of WT viruses after infection for 7 hours. In the presence of low pH-inhibitors, the fluorescence intensity of DiO in cells infected by WT and mutant viruses reduced at the same degree (S9C Fig). These results indicated that H144A and Q258A substitutions reduced the membrane fusion activity of JEV. Taken together, H144 and Q258 are dispensable for the intracellular transport of JEV but essential for the membrane fusion with endosomes.

## Conclusion

We developed a lipid-specific method to mildly, readily, specifically, and efficiently label enveloped viruses with QDs. The unique optical properties of QDs, the high specificity and efficiency, and the comparative convenience make it superior to the DiD and DiO labeling. The advantages in convenience and universality make this lipid-specific method prevail over other QD labeling methods. More importantly, since the target molecules are lipids, this method is competent to label key protein-mutated viruses, which is significant for in-depth study of virus infection mechanisms. And because this labeling method does not involve virus propagation, it can also be used to study inactivated high virulent viruses such as HIV and Ebola virus. The labeling technique described in this study provides a powerful tool to visually investigate the dynamic infection of enveloped viruses.

## Materials and methods

### Cells

Vero, Madin-Darby canine kidney (MDCK), Baby hamster kidney (BHK-21), and MARC-145 cells were maintained in Dulbecco’s modified Eagle’s medium (DMEM, Gibco) supplemented with 10% fetal bovine serum (FBS, South America origin, PAN Biotech) under 5% CO_2_ at 37°C.

### Viruses

JEV SA-14-14-2 and WT JEV AT31 were propagated in BHK-21 cells. H144A and Q258A mutant JEV virions were packaged using cDNA clones of JEV AT31 as described [35]. PRRSV (HN07-1 strain) was propagated in MARC-145 cells. Collected JEV- and PRRSV-containing cell culture supernatant was centrifuged at 1500 rpm and 4°C for 10 min and filtered with 0.2 μm pore size filters (Millipore) to remove cell debris. To evaluate the effect of ultracentrifugation on virus infectivity, part of the JEV SA-14-14-2 was further purified by ultracentrifugation [38, 39]. In brief, viruses were concentrated by centrifugation at 100,000 × *g* and 4°C in a Ty45 Ti rotor (Beckman) for 2 hours, then purified by gradient centrifugation on 10-35% potassium tartrate-glycerol (30%) at 125,000 × *g* in a SW32 Ti rotor for 2 hours, and desalted at 180,000 × *g* in the Ty45 Ti rotor for 1 hour. IAV (A/chicken/Hubei/01-MA01/1999(H9N2) strain) was propagated in pathogen-free chicken eggs and purified by sucrose gradient ultracentrifugation as described [40]. All the harvested viruses were subpackaged and stored at −80°C until use.

### Virus labeling

Viruses were incubated with 30 μM DSPE-PEG (2000)-Biotin (Avanti) at room temperature for 1 hour. Unincorporated biotin and aggregated viruses were removed by NAP-5 gel filtration columns (GE Healthcare) and 0.2 μm pore size filters, respectively. Then, biotinylated viruses and 2 nM SA-QD 705 (Wuhan Jiayuan Quantum Dots Co., Ltd.) were successively incubated with cells at 4°C for 30 and 10 min, respectively, allowing viruses to pre-bind to cell surfaces and QDs to bind to viruses. Unbound viruses and QDs were removed by washing cells with ice-cold PBS. To track virus infection, the cells were immediately warmed to 37°C and imaged on a spinning-disk confocal microscope equipped with a cell culture system.

Labeling of viruses with DiD/DiO was done by incubating viruses with 5 μM DiD/DiO (Beyotime Biotechnology) under shaking and in the dark at room temperature for 1 hour. Labeling of viruses with both DiO and R18 was done by incubating viruses with 0.2 μM DiO and 0.4 μM R18 (Millipore) under the same conditions. Unbound dyes and aggregates were removed by gel filtration and syringe filtration.

### Immunofluorescence assay

Anti-Japanese encephalitis E (mouse monoclonal, Millipore), influenza A H9N2 HA (mouse monoclonal, Sino Biological Inc.), and PRRSV nucleocapsid protein (rabbit monoclonal, VMRD) antibodies were used to localize JEV. IAV, and PRRSV, respectively. DyLight 488/649 conjugated secondary antibodies (Abbkine) were used to label the primary antibodies, illuminating the viruses.

### Virus infectivity

The infectious infectivity of JEV and the number of GCPs were measured by plaque assay on BHK-21 cells and quantitative PCR (qPCR) as described [35]. PRRSV infectivity was measured by TCID_50_ on Vero cells. IAV infectivity was measured by TCID_50_ assay on MDCK cells and hemagglutination assay on red blood cells (41). NHS-Biotin-IAV was obtained as described (40). Briefly, 100 μL of IAV was incubated with 0.1 mg Sulfo-NHS-LC-Biotin (Thermo) at room temperature for 2 hours. Unbound biotin and aggregates were removed by filtration.

### TEM imaging

Twenty μL of 10 nM SA-QD 705, JEV, and biotinylated JEV incubated with 0.1 nM SA-QD 705 were dropped on carbon-coated copper grids, respectively. After 0.5 hour (for SA-QD 705) or 15 hours (for JEV and QD-labeled JEV) at 4°C, the grids were drained by filter papers and washed with ultrapure water. After being stained with sodium phosphotungstate for 3 min (for JEV) or 30 s (for QD-labeled JEV), the grids were air dried and imaged on a HITACHI-7000FA transmission electron microscope.

### Fluorescence imaging

Fluorescence images were captured by a spinning-disk confocal microscope (Andor Revolution XD). Hoechst 33342, DyLight 488/DiO, R18, and Dylight 649/DiD/CellMask deep red plasma membrane stain were imaged using 405, 488, 561, and 640 nm lasers (DPSS Lasers Inc.) and 447/60, 525/50, 605/20, and 685/40 nm emission filters (Chroma), respectively. QD 605 and QD 705 were imaged using the 488 nm laser and 605/20 and 685/40 nm emission filters.

### Image analysis

Colocalization events were evaluated by Mander’s coefficient and intensity correlation analysis (ICA) using ImageJ [31, 42]. Mander’s coefficients vary from 0 (non-overlapping images) to 1 (100% colocalized images) and are termed as tM_QD_ and tM_DyLight_ here according to the image names. tM_QD_ is the ratio of the ‘summed intensities of the QD signal colocalized with DyLight signals’ to the ‘total intensities of QD signals’ in thresholded images, and tM_DyLight_ is defined conversely. ICA is based on the assumption that the summed difference of pixel intensities from the mean in a single channel is zero, namely ∑_n pixels_ (*I*_QD, i_ − *I*_QD, mean_) = 0 and ∑_n pixels_ (*I*_DyLight, i_ − *I*_DyLight, mean_) = 0. PDM is the product (*I*_QD, i_ − *I*_QD, mean_)(*I*_DyLight, i_ − *I*_DyLight, mean_). Intensity correlation plots show the intensity as a function of PDM. ICQ is the ratio of the ‘summed positive PDM from two channels’ to the ‘total PDM’ subtracted by 0.5. It varies from –0.5 (mutual exclusion) to +0.5 (complete colocalization) and indicates a strong covariance in the range from 0.1 to 0.5 [43]. Line profiles of signals were acquired with Image-Pro Plus.

Trajectories of viruses were reconstructed by linking points in each frame using the nearest-neighbor association and the motion history of individual particles with Image-Pro Plus [44, 45]. MSD representing the average squared distance of all steps within a trajectory for *Δt* (*Δt* = *τ*, 2*τ*, 3*τ*, and so on, *τ* = acquisition time interval between frames) was calculated using MATLAB [46]. Modes of motion were analyzed by fitting MSD and *Δt* to functions: MSD = 4*DΔt* (normal or Brownian diffusion), MSD = 4*DΔt* + (*VΔt*)^2^ (active or directed diffusion), and MSD = 4*DΔt*^*α*^ (anomalous diffusion) [47].

### Statistical analysis

Data are represented as mean ± SD. Student’s *t-*test was performed for all statistical analyses. Statistical significance was determined by two-tailed P values: ns P >0.05, *P <0.05, **P <0.01, ***P <0.001.

## Acknowledgements

We thank Cai-Ping Wang and Peng-Juan Li from Henan Agricultural University for providing PRRSV and MARC-145 cells.

## Supporting information

**S1 Fig. Labeling cell membranes with QDs by the rapid insertion of lipid-biotin conjugates into membranes.** (A) Cells incubated with 5 μM DiD (red) for 0, 30, 60, 120, 180, and 240 min were imaged by a confocal microscope. (B) Cells were incubated with 30 μM DSPE-PEG-Biotin for 0, 30, 60, 120, 180, and 240 min and then with SA-modified QDs for 10 min (red). CellMask Deep Red Plasma Membrane Stain (green) and Hoechst 33342 (blue) were used to stain the plasma membrane and the nucleus. Scale bars, 10 μm. (C, D) Mean fluorescence intensity (MFI) of the DiD/QD-labeled cells and the labeling efficiency measured by FCM (n=3).

**S2 Fig. Specifically and efficiently labeling JEV with QDs on cell surfaces.** (A) JEV and biotinylated JEV were pre-attached to cell surfaces and labeled with SA-QDs (red) at 4°C. After fixation, viruses on cell surfaces were further labeled with anti-E-DyLight 488(green). Scale bars, 10 μm. (B) Line profile showing distributions of the QD and DyLight signals on the line in A. (C) PDM image showing the colocalized and uncolocalized spots in the lower merge panel in A. (D) The tM_QD_, tM_DyLight_, and ICQ values calculated from 30 randomly selected cells.

**S3 Fig. Low specificity and efficiency of DiD and DiO labeling.** (A, D) DiD/DiO-labeled JEV (red) were attached to Vero cell surfaces at 4°C and further labeled with anti-E-DyLight 488/649 (green) after fixation. Images were acquired by the same confocal microscope setup. Scale bars, 10 μm. (B, E) Signals in the overlapped images in A and D were randomly connected with lines. The line profiles show distributions of DiD/DiO and DyLight signals on the lines. (C, F) The tM_DiD/DiO_, tM_DyLight_, and ICQ values calculated from 40 randomly selected cells.

**S4 Fig. Quantification of DSPE-PEG-Biotin on single biotinylated JEV.** (A) Schematic representation of the method used to quantify biotin with SA-FITC conjugates. (B) Fluorescence spectra of SA-FITC solution titrated with biotin. Lines from the bottom to the top show the biotin consumption of 0, 0.5, 1.0, 1.5, 2.0, 2.5, 3.0, 3.5, 4.0, 4.5, and 5.0 pmol. (C) The fluorescence intensity of SA-FITC at 515 nm. (D) The dependence between fluorescence intensity of SA-FITC and the biotin consumption (symbols). The red line is the fit to y = 209677x + 238115. The fluorescence intensity of SA-FITC has good linear relation with biotin consumption in the range from 0.5 to 2.5 pmol. (E) Fluorescence spectra of SA-FITC solution added with 0.5 pmol biotin, 5 μL of JEV, and 5 μL of biotinylated JEV. (F) The amount of DSPE-PEG-Biotin on single JEV virions (n=3).

**S5 Fig. Quantification of QD 705 on single JEV.** (A) Fluorescence spectra of QD 705 and QD-labeled JEV. This data indicated that combining with viruses didn’t change QD fluorescence. (B) Statistic gray levels of QD 705 subtracted by that of the noise. The red line is the fit to Gaussian function and the mean is 449.8, indicating that the gray level of most QD particles is about 450. (C) The trace of a QD with gray level of about 450 (left) and a zoom of it (right), showing that this QD particle has obvious blinking behaviors and is a single QD. (D) About 95% QDs with gray levels of about 450 are single QD (n=113). Results from B-D suggest that the gray level of single QD is around 450. (E) Statistic gray lavels of QD-labeled JEV subtracted by that of brackground, showing that the gray level of most virions is around 900 and 1350. That means most JEV virions combined with 2 or 3 QDs.

**S6 Fig. Specifically and efficiently labeling PRRSV with QDs.** (A) PRRSV was biotinylated with DSPE-PEG-Biotin, attached to Vero cell surfaces, and then labeled with SA-QD 705 (red). After fixation, PRRSV was further labeled with anti-PRRSV-DyLight 488 (green). (B) Intensity correlation plots (ICPs) of the QD and DyLight signals in A and their scatter plot. The C-shaped curves of dots in ICPs and the centred dots in the scatter plot show that QD and DyLight signals are almost completely colocalized. (C) PDM image of the double-labeled viruses showing the colocalized and uncolocalized spots. (D) Line profile showing distributions of the QD and DyLight signals on the line in the overlapped image of A.

**S7 Fig. Specifically and efficiently labeling IAV with QDs.** (A) IAV was biotinylated with DSPE-PEG-Biotin, attached to MDCK cell surfaces, and labeled with SA-QD 705 (red). After fixation, IAV was further labeled with anti-HA-DyLight 488 (green). (B) ICPs of QD and Dylight signals and the scatter plot. (C) PDM image showing the colocalized and uncolocalized spots in the overlapped image of A. (D) Line profile showing distributions of the signals on the line in A.

**S8 Fig. JEV transport *via* a microfilament-independent and microtubule/dynein-dependent pathway.**

(A) QD-labeled JEV was allowed to infect Vero cells treated with 0.2% DMSO, 20 μM cytochalasin D (CytoD), 60 μM nocozadole (Noc), and 100 μM ciliobrevin D (CilioD). CytoD, Noc, and CilioD were used to block microfilaments, microtubules, and dynein, respectively. After 0.5 h of virus uptake, viruses remained on cell surfaces were stained with SA-Cy3 at 4°C to be distingushed from the internalized viruses. After fixation, the cells were imaged in 3D and analyzed with Velocity. Horizontal scale bars, 10 μm. Vertical scale bars, 5 μm. (B) The amount of viruses internalized in cells treated with drugs (n=50). (C) Virus infection in cells treated with drugs were tracked. The white lines are trajectories of viruses. (D) The speed *vs.* time plots of the viruses tracked in C. (E) The MSD *vs. Δt* plots (black symbols). The upward lines in the first two graphs are the fits to MSD = 4*DΔt* + (*VΔt*)^2^ with D = 0.031/0.014 μm^2^/s and V = 0.088/0.10 μm/s. The downward line in the third graph is the fit to MSD = 4*DΔt*^*α*^ with D = 0.0012 μm^2^/s and α = 0.77.

**S9 Fig. H144A and Q258A substitutions inhibiting JEV fusion with endosomes.** (A) DiO/R18 double-labeled viruses were allowed to infect Vero cells for different times. Scale bar, 10 μm. (B) MFI of DiO in cells infected by the double-labeled viruses for different times measured by FCM. (C) MFI of DiO in cells treated by drugs and infected by JEV for 1 h. NH_4_Cl and chloroquine (CQ) were used to block the virus-endosome fusion.

**S1 Movie. WT JEV moving from cell periphery toward the cell interior.** WT JEV (red) was labeled with QD 705 and imaged with frame intervals of 0.63 s for about 100 s.

**S2 Movie. H144A JEV moving from the cell periphery toward the cell interior.** H144A JEV (red) was labeled with QD 705 and imaged with frame intervals of 0.73 s for about 191 s.

**S3 Movie. Q258A JEV moving from the cell periphery toward the cell interior.** Q258A JEV (red) was labeled with QD 705 and imaged with frame intervals of 0.23 s for about 41 s.

